# Novel methods to identify TCM constitution of hyperlipidemic patients and predict atherosclerotic diseases

**DOI:** 10.1101/2022.07.20.500876

**Authors:** Liling Zeng, Qixin Zhang, Chen Sun, Li Huang, Jiamin Yuan, Fei Tan, Yanhua Wu, Zhimin Yang, Fuping Xu

**Affiliations:** Department of Traditional Chinese Medicine, Guangzhou Red Cross Hospital, Medical College, Jinan University, Guangzhou 510000, China; ZILKHA Neurogenetic Institute, Keck School of Medicine of USC, Los Angeles 90001, United State; The Second Affiliated Hospital of Guangzhou University of Chinese Medicine, Guangzhou 510000, China

**Keywords:** Differentially expressed microRNAs, TCM constitution, hyperlipidemia, predisposition, atherosclerotic diseases

## Abstract

Hyperlipidemia can accelerate the progression of atherosclerosis, ultimately leading to cardiovascular disease. TCM constitution has been widely used as an indicator of health status and susceptibility to diseases. However, there still lack of objective, credible ways to identify TCM constitution of hyperlipidemic patients, and the connection between TCM constitution and atherosclerotic diseases in individuals with hyperlipidemia is unknown. This study aims to identify differentially expressed microRNAs (DEMs) as biomarkers of the TCM constitution of hyperlipidemic patients and explore the potential connection between TCM constitution and atherosclerotic diseases. In the study, we randomly recruited 10 hyperlipidemic patients with asthenic constitution (HAC), 10 hyperlipidemic patients with strong constitution (HSC), and 10 hyperlipidemic patients with normal constitution (HNC) and collected serum miRNA. After analyzing the miRNA expression profiles, we found that hsa-miR-338-3p may be a potential blood biomarker for the identification of the TCM constitution of hyperlipidemic patients. Moreover, the HSC classification is likely a cardiocerebrovascular disease predisposition and is closely related to the inflammatory process and glucose and lipid metabolism, which contribute to the development of atherosclerotic diseases.

## Introduction

Hyperlipidemia, especially hypercholesterolemia and high levels of low-density lipoprotein cholesterol (LDL-C), accelerates atherosclerotic plaque formation in patients with cerebrovascular disease^[1]^or coronary heart disease (CHD)^[2]^. These are diseases that have emerged as the most frequent cause of death and major disability worldwide^[3, 4]^. At present, lipid-lowering therapy is one of the main interventions for CHD and cerebrovascular disease^[5–7]^. Currently, statins, such as lovastatin and atorvastatin, are the main therapies to lower LDL-C^[8, 9]^but have a number of adverse effects, including muscle myopathy and derangements in hepatic function^[10,11]^. Therefore, novel preventive and therapeutic strategies for hyperlipidemia still need to be developed.

In Eastern countries, in addition to Western medical treatment, traditional Chinese medicine (TCM) is highly accessible and widely used in patients with hyperlipidemia. Numerous studies have shown that TCM therapy could have lipid-lowering and lipid-controlling effects in patients with hyperlipidemia^[12–14]^. TCM constitution can be used as an indicator of health status and susceptibility to diseases^[15]^. In addition, according to the theory of differentiation syndrome and treatment with Chinese medicine, treatments used in hyperlipidemic patients should be based on the TCM constitution of patients. Moreover, the TCM constitution of hyperlipidemic patients is also a predisposing factor for atherosclerosis and cardiovascular disease, which is a common concomitant disease in individuals with hyperlipidemia. Previous studies^[16]^ revealed that TCM constitution has the potential to be a first-line diagnostic tool to facilitate early recognition and diagnosis of CAD. A systematic review also studied the correlation between TCM constitution and dyslipidemia and provided evidence for the prevention and treatment of dyslipidemia using TCM^[17]^. Thus, identification of the TCM constitution of hyperlipidemic patients is very important for treating hyperlipidemia and preventing the development of atherosclerosis and cardiovascular disease. However, the main method to identify TCM constitution of hyperlipidemic patients is based on doctors’ experience, and there still lack of objective, credible ways to assess TCM constitution.

MicroRNAs (miRNAs), a type of noncoding RNA, have attracted attention, as they have been applied in multiple fields of life science and medicine. In previous decades, microarray analysis has been widely used to investigate differentially expressed miRNAs (DEMs), dysregulated genes and pathways associated with TCM constitution^[18]^. However, there is little evidence that identifies DEMs associated with the TCM constitution of hyperlipidemic patients, and there is no research that has explored the potential connection between TCM constitution and the development of atherosclerotic diseases in individuals with hyperlipidemia. Thus, in the present study, expression data obtained from hyperlipidemic patients with different TCM constitutions were subjected to a bioinformatics analysis to provide more reliable data. In this study, we applied DEMs as biomarkers to identify the TCM constitution of hyperlipidemic patients and explored the underlying risk factors involved in the development of atherosclerosis and cardiovascular disease in individuals with hyperlipidemia.

## Methods

### Study Subjects and Sampling Process

The present study was approved by the Research Ethics Committees of Guangdong Provincial Hospital of TCM in accordance with the Declaration of Helsinki, and written informed consent was obtained from each participant. All patients were selected with standard diagnostic codes and exclusion and inclusion rules from Sep. 8th, 2017 to Sep. 20th, 2018. The details about the diagnosis standard, exclusion and inclusion criteria are shown in Table 1. Hyperlipidemia was diagnosed according to the Guidelines^[19]^ for the prevention and treatment of dyslipidemia in Chinese adults and was defined as fasting plasma levels of TC ≥ 6.2 mmol/L (240 mg/dl), TG ≥ 2.3 mmol/L (200 mg/dl), HDL-C ≥ 1.0 mmol/L (40 mg/dl), LDL-C ≥ 4.1 mmol/L (160 mg/dl), or non-HDL-C ≥ 4.9 mmol/L (190 mg/dl). The identification of TCM constitution was performed according to the Guidelines for the classification and recognition of TCM constitution (ZYYXH/T157-2009)^[20]^. In the present study, TCM constitutions in hyperlipidemia were classified into three types: hyperlipidemic patients with asthenic constitution (HAC, namely, Xuzheng); hyperlipidemic patients with normal constitution (HNC, namely, Pinghezhi); and hyperlipidemic patients with strong constitution (HSC, namely, Shizheng). At the screening stage, a total of 1041 subjects were screened, and 347 hyperlipidemic patients (234 HNC, 53 HSC and 60 HAC) were included (Figure 1). Finally, a total of 30 subjects, 10 HAC, 10 HSC and 10 HNC, were randomly assigned to groups in this study. All subjects were recruited from the village community at Foshan, Guangdong, China, from September 2017 to December 2018. All subjects provided a detailed history and underwent physical examination and blood tests. Full blood samples were obtained from participants at least 12 h following their most recent meal. The blood samples were collected in serum vacutainer tubes with a clot activator, and blood serum was immediately separated by a 10-min centrifugation at 1500 ×g. Clinical biochemical parameters such as total cholesterol (TC), were measured. The remaining serum samples were stored at −80°C until miRNA analysis.

**Figure 1.**
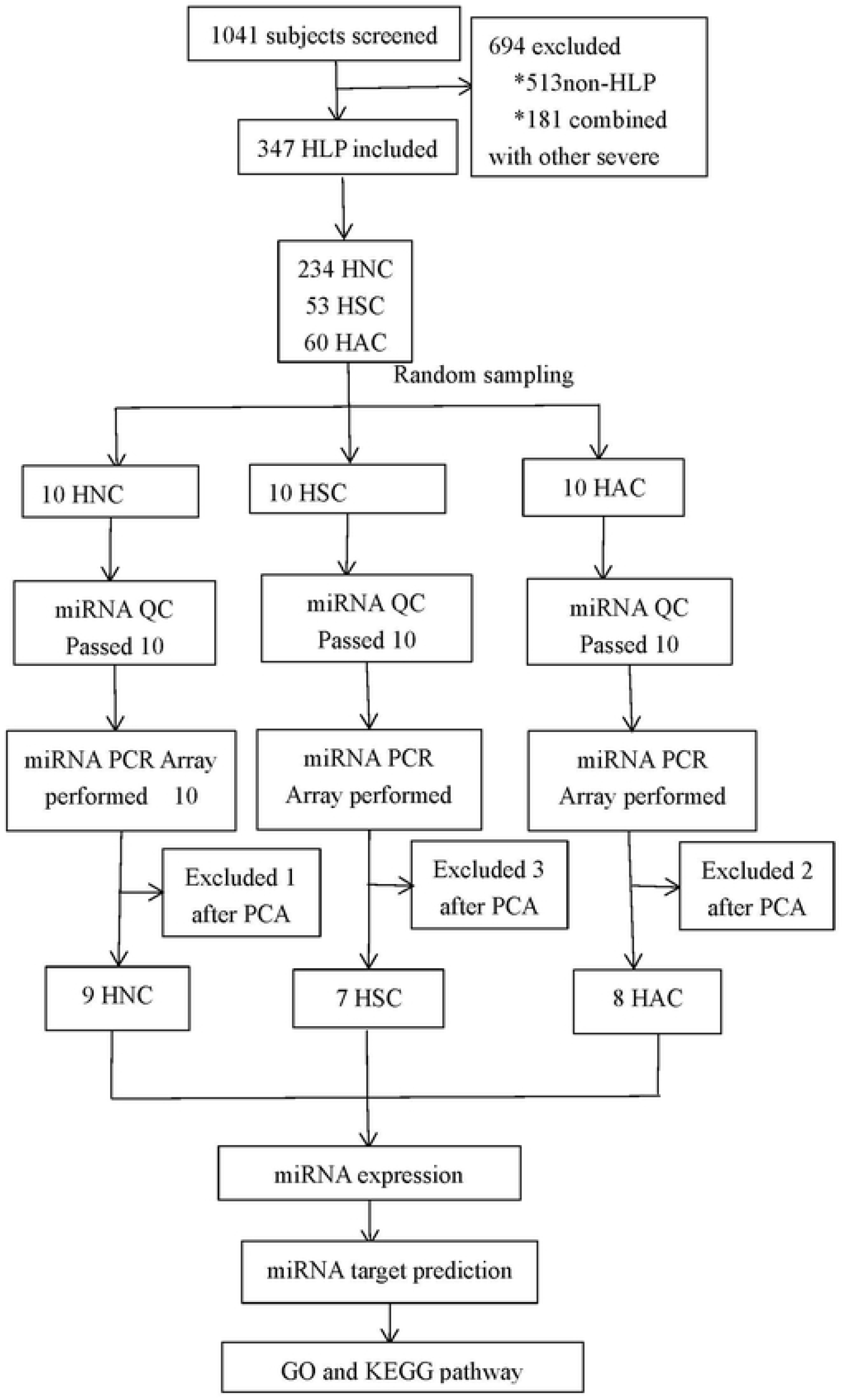
Study Flow Diagram HLP, hyperlipidemia; HAC, hyperlipidemic patients with asthenic constitution; HNC, hyperlipidemic patients with normal constitution; HSC, hyperlipidemic patients with strong constitution; PCA, principal component analysis; QC, quality control; GO, Gene Ontology; KEGG, Kyoto Encyclopedia of Genes and Genomes.

**Table 1.**
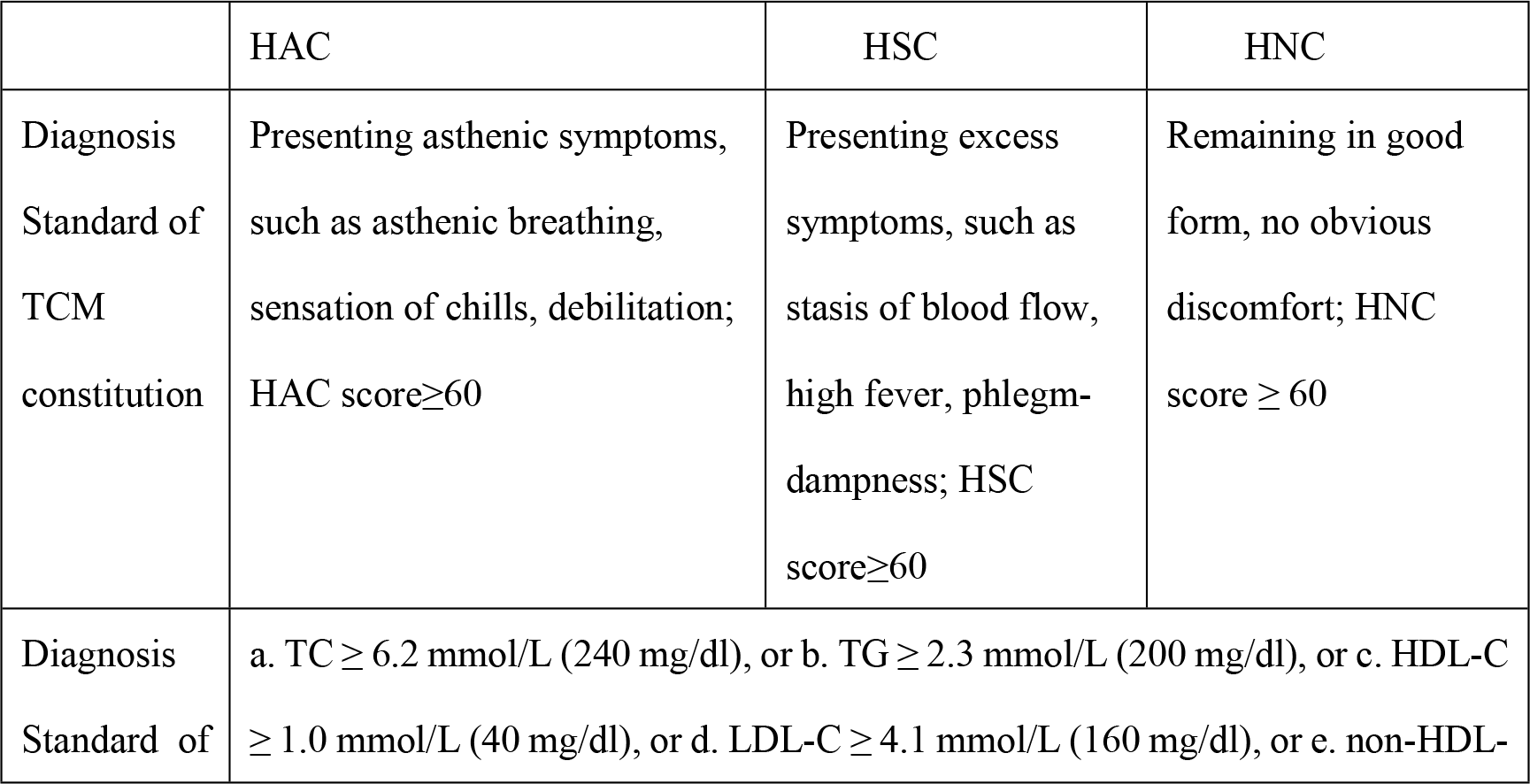
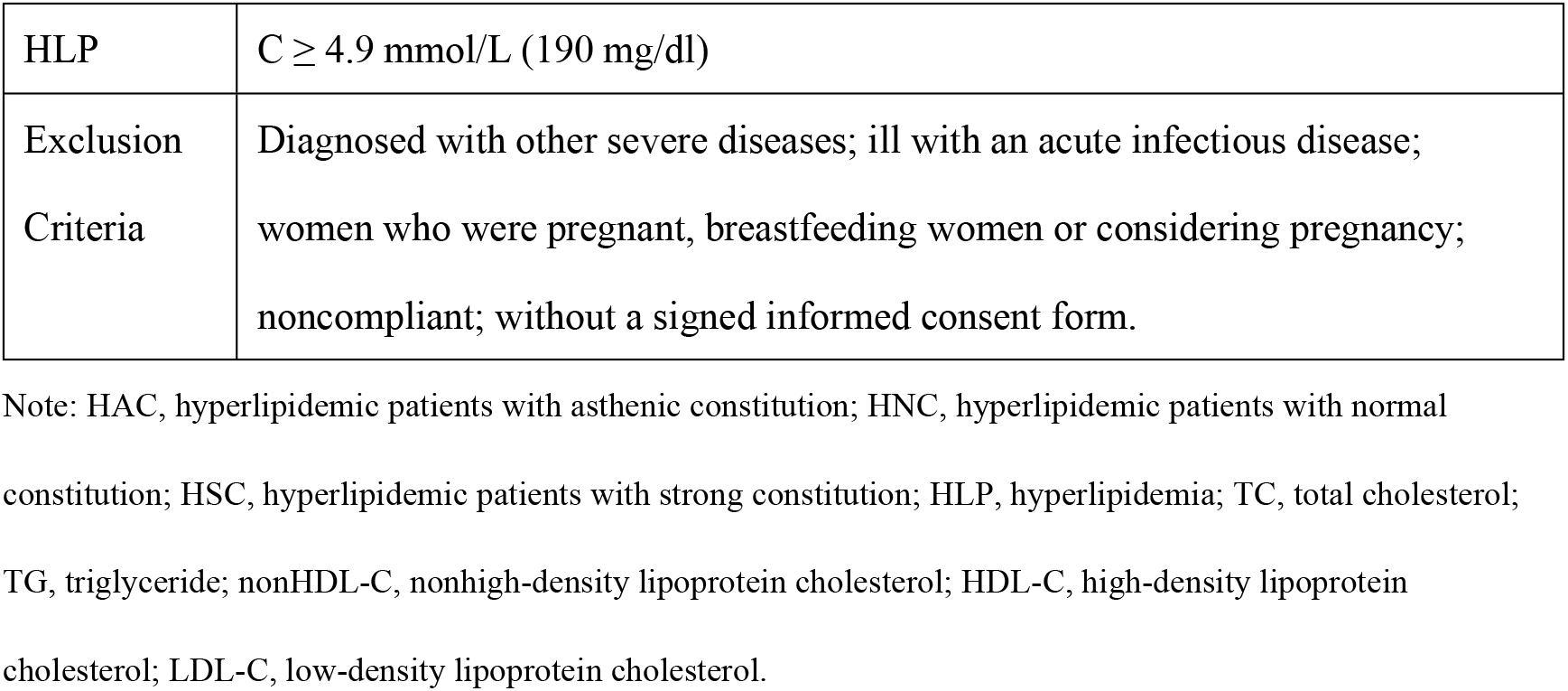
Diagnosis Standard, Inclusion and Exclusion Criteria

|  | HAC | HSC | HNC |
| --- | --- | --- | --- |
| Diagnosis Standard of TCM constitution | Presenting asthenic symptoms, such as asthenic breathing, sensation of chills, debilitation; HAC score≥60 | Presenting excess symptoms, such as stasis of blood flow, high fever, phlegm-dampness; HSC score≥60 | Remaining in good form, no obvious discomfort; HNC score ≥ 60 |
| Diagnosis Standard of | a. TC ≥ 6.2 mmol/L (240 mg/dl), or b. TG ≥ 2.3 mmol/L (200 mg/dl), or c. HDL-C ≥ 1.0 mmol/L (40 mg/dl), or d. LDL-C ≥ 4.1 mmol/L (160 mg/dl), or e. non-HDL- | | |

|  |  |
| --- | --- |
| HLP | $C \geq 4.9$ mmol/L (190 mg/dl) |
| Exclusion<br>Criteria | Diagnosed with other severe diseases; ill with an acute infectious disease;<br>women who were pregnant, breastfeeding women or considering pregnancy;<br>noncompliant; without a signed informed consent form. |

Note: HAC, hyperlipidemic patients with asthenic constitution; HNC, hyperlipidemic patients with normal constitution; HSC, hyperlipidemic patients with strong constitution; HLP, hyperlipidemia; TC, total cholesterol; TG, triglyceride; nonHDL-C, nonhigh-density lipoprotein cholesterol; HDL-C, high-density lipoprotein cholesterol; LDL-C, low-density lipoprotein cholesterol.

### Microarray Information

The QIAGEN miScript PCR System (QIAGEN, Germany) was used in this experiment. The miScript PCR System consists of the miScript II RT Kit, miScript PreAMP PCR Kit, miScript miRNA PCR Array, miScript Primer Assay, miScript SYBR Green PCR Kit, and miScript miRNA PCR Array data analysis tool. The miScript PCR System allows sensitive and specific detection and quantification of miRNAs. The miScript PCR System uses total RNA that contains miRNA as the starting material for cDNA synthesis, and separate enrichment of small RNA is not needed. The miScript miRNA PCR Arrays consist of mature miRNA-specific forward primers (miScript Primer Assays) that have been arrayed in biologically relevant pathway-focused and whole miRNome panels. These PCR arrays are provided in ready-to-use 384-well plates. Each assay in a miScript miRNA PCR Array has been verified to ensure sensitive and specific detection of mature miRNAs by real-time PCR. The free, web-based miScript miRNA PCR Array data analysis tool (https://www.qiagen.com/cn/shop/genes-and-pathways/data-analy) simplifies the analysis of real-time PCR data. Once raw threshold cycle (CT) data have been uploaded, the tool automatically performs all fold change calculations using the ΔΔCT method of relative quantification and presents the results in several formats. Mature miRNome expression profiling is now within the reach of every laboratory because of the ease, convenience, and consistent performance of the miScript miRNA PCR Arrays. The miScript miRNA PCR arrays are at the forefront of real-time PCR-based mature miRNA profiling tools.

### Experimental Design

The miRNeasy Serum/Plasma Kit (QIAGEN, Cat# 217184) was used for the extraction and purification of total RNA from human serum. The miRNeasy Serum/Plasma Spike-In Control was used for the normalization of miRNAs purified from human serum. After the purification of total RNA, which contains mature miRNAs, total RNA was quantified by a NanoDrop ND-2000 (Thermo Scientific). The RNA integrity was assessed using an Agilent Bioanalyzer 2100 (Agilent Technologies).

The expression profile of miRNAs was tested based on the manufacturer’s standard protocols by using the miScript PCR System (QIAGEN), which consists of the miScript II RT Kit, miScript PreAMP PCR Kit, miScript miRNA PCR Array, miScript Primer Assay, miScript SYBR Green PCR Kit, and miScript miRNA PCR Array data analysis tool. Briefly, total RNA was reverse transcribed to double-stranded cDNA after incubation at 37°C for 60 min and then at 95°C for 5 min. Then, the PCR mixture was prepared and added to the miScript miRNA PCR Array. Finally, real-time PCR was performed, and the results were analyzed using the miScript miRNA PCR Array data analysis tool.

### Data Analysis

Clinical data were analyzed with SPSS 18.0 software (version 18; SPSS Inc., Chicago, IL, USA). Fisher’s exact test was used to compare the categorical variables, and oneway ANOVA was used to compare the continuous variables among groups. The data are presented as the mean and standard error of the mean (SEM) for continuous variables and absolute numbers and percentages for categorical variables.

For the analysis of miRNA expression profiles, the miScript miRNA PCR Array data analysis tool (QIAGEN 5.3) was used to analyze the raw data. First, the raw data were normalized to the median. DEMs were then identified based on a fold change and *P*-values were calculated with t tests. The threshold set for up- and downregulated genes was fold change ≥2 or ≤0.5 and *p*-value <0.05. Afterward, Gene Ontology (GO) and Kyoto Encyclopedia of Genes and Genomes (KEGG) analyses were applied to determine the roles of mRNAs associated with the DEMs. A Venn diagram was used to identify the overlapping DEMs among the groups.

## Results

After excluding the outliers identified by PCA (principal component analysis) (Figure S1), there were 9 samples in HNC, 7 in HSC, and 8 in HAC whose data were used for the final analysis.

### Clinical Characteristics of Study Subjects

When the patients were recruited, there were no significant differences in the distribution of age, sex, or BMI distribution among the three study groups (*p* >0.05, Table 2). However, among the three groups, HSC group was more likely to have diabetes and hyperuricemia and had higher levels of FT3 and uric acid (*p* <0.05, Table 2), and HAC group had lower red blood cell counts and hemoglobin levels (*p* <0.05, Table 2).

**Table 2.** Characteristics of the study patients

|  | HNC (n=9) | HSC (n=7) | HAC (n=8) | *P* value |
| --- | --- | --- | --- | --- |
| Age (mean ± SD) | 51.44±7.06 | 46.57±4.34 | 59±7.21 | 0.440 |
| Sex (n (%)) |  |  |  |  |
| Female | 5 (55.6) | 4 (57.1) | 7 (87.5) | 0.309 |
| Male | 4 (44.4) | 3 (42.9) | 1 (12.5) |  |
| BMI (kg/m<sup>2</sup>, mean ± SD) | 22.80±1.21 | 24.31±1.37 | 21.64±1.09 | 0.345 |
| Smoking (n (%)) |  |  |  |  |
| No | 8 (88.9) | 5 (71.4) | 7 (87.5) | 0.277 |
| Yes | 1 (11.1) | 2 (28.6) | 1 (12.5) |  |
| Drinking (n (%)) |  |  |  |  |
| No | 7 (77.8) | 5 (71.4) | 7 (87.5) | 0.328 |
| Yes | 2 (22.2) | 2 (28.6) | 1 (12.5) |  |
| Diabetes (%) |  |  |  |  |
| No | 9 (100) | 4 (57.1) | 8 (100) | 0.014 |
| Yes | 0 (0) | 3 (42.9) | 0 (0) |  |
| Hyperuricemia (%) |  |  |  |  |
| No | 7 (77.8) | 3 (42.9) | 8 (100) | 0.019 |
| Yes | 2 (22.2) | 4 (57.1) | 0 (0) |  |
| Chronic inflammatory diseases |  |  |  |  |
| No | 7 (77.8) | 5 (71.4) | 6 (75) | 0.959 |
| Yes | 2 (22.2) | 2 (28.6) | 2 (25) |  |
| Chronic renal failure |  |  |  |  |
| No | 5 (55.6) | 4 (57.1) | 7 (87.5) | 0.273 |
| Yes | 4 (44.4) | 3 (42.9) | 1 (12.5) |  |
| Anemia |  |  |  |  |
| No | 9 (100) | 7 (100) | 6 (75) | 0.092 |
| Yes | 0 (0) | 0 (0) | 2 (25) |  |
| FT3 (mean ± SD) | 5.51±0.11 | 5.61±0.13 | 4.89±0.19 | 0.006 |
| FT4 (mean ± SD) | 17.15±0.15 | 16.96±0.77 | 16.41±2.07 | 0.920 |
| PCT (mean ± SD) | 0.28±0.01 | 0.26±0.01 | 0.28±0.02 | 0.670 |
| WBC (mean ± SD) | 8.02±0.37 | 6.63±0.32 | 6.16±0.58 | 0.016 |
| RBC (mean ± SD) | 4.96±0.15 | 4.86±0.15 | 4.23±0.22 | 0.014 |
| PLT (mean ± SD) | 265.78±16.47 | 247.57±14.97 | 266.13±22.59 | 0.740 |
| TSH (mean ± SD) | 1.67±0.29 | 2.29±0.42 | 2.63±1.38 | 0.706 |
| TSTO (mean ± SD) | 9.96±3.11 | 7.86±3.33 | 3±1.58 | 0.237 |
| Hs-CRP (mean ± SD) | 2.46±0.86 | 3.97±1.37 | 5.99±2.86 | 0.045 |
| Serum creatinine (mean ± SD) | 77.22±5.7 | 68.42±6.84 | 62.62±3.40 | 0.169 |
| eGFR (mean ± SD) | 72.19±14.85 | 61.27±22.43 | 83.46±13.12 | 0.666 |
| Hb (mean ± SD) | 144.33±4.18 | 150.14±5.8 | 129.50±5.52 | 0.030 |
| UA (mean ± SD) | 350.67±32.57 | 426.71±29.98 | 292±16.96 | 0.012 |
| GLU (mean ± SD) | 3.99±0.07 | 5.3±0.78 | 5.43±0.64 | 0.125 |
| TC (mean ± SD) | 6.12±0.38 | 6.48±0.39 | 6.79±1.23 | 0.823 |
| TG (mean ± SD) | 1.12±0.09 | 2.14±0.59 | 1.3±0.18 | 0.087 |
| NonHDL-C (mean ± SD) | 4.48±0.33 | 4.83±0.42 | 5.18±1 | 0.744 |
| HDL-C (mean ± SD) | 1.64±0.12 | 1.65±0.20 | 1.62±0.24 | 0.991 |
| LDL-C (mean ± SD) | 4.07±0.24 | 4.1±0.27 | 4.3±0.55 | 0.899 |

**Note:** BMI, body mass index; FT3, free triiodothyronine; FT4, free thyroxine; PCT, procalcitonin; WBC, white blood cell count; RBC, red blood cell count; PLT, platelet count; TSH, thyroid stimulating hormone; TSTO, testosterone; Hs-CRP, high-sensitivity C-reactive protein; eGFR, glomerular filtration rate; Hb, hemoglobin; UA, uric acid; GLU, glucose; TC, total cholesterol; TG, triglyceride; nonHDL-C, nonhigh-density lipoprotein cholesterol; HDL-C, high-density lipoprotein cholesterol; LDL-C, low-density lipoprotein cholesterol.

### Microarray Data Analysis of DEMs Among the Three Groups

To identify DEMs after pairwise comparisons, fold change ≥2 or ≤0.5 and *p* value <0.05 were set as the cut-off criteria when screening the miRNA array profiles. Compared to the HNC group, 25 downregulated miRNAs were identified to be dysregulated in the HSC group (Table S1, Figure S2), and 8 miRNAs (7 upregulated and 1 downregulated) were identified to be dysregulated in the HAC group (Table S2, Figure S3). Six miRNAs were identified to be upregulated in the HSC group compared to the HAC group (Table S3, Figure S4).

To identify the common DEMs among the three groups, Venn diagram analysis was performed (Figure 2). The results showed that hsa-miR-338-3p was the common DEM among the groups; it was downregulated in HSC and upregulated in HAC (Figure 3). Below, we summarize the characteristics of all the DEMs found in the three groups (Table 3).

**Figure 2.**
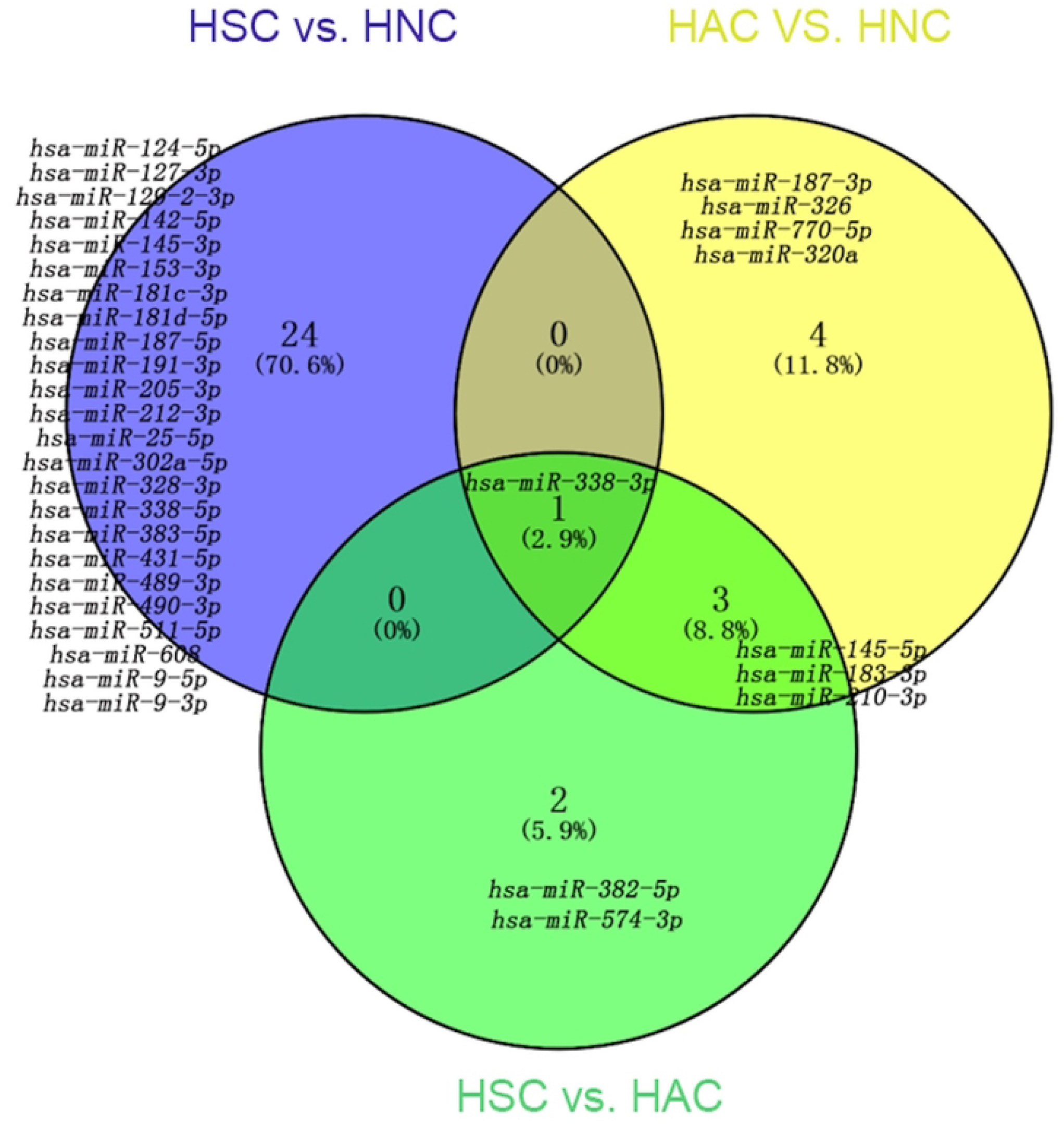
Venn diagram. Hsa-miR-338-3p was identified as the common DEM among the groups.

**Figure 3.**
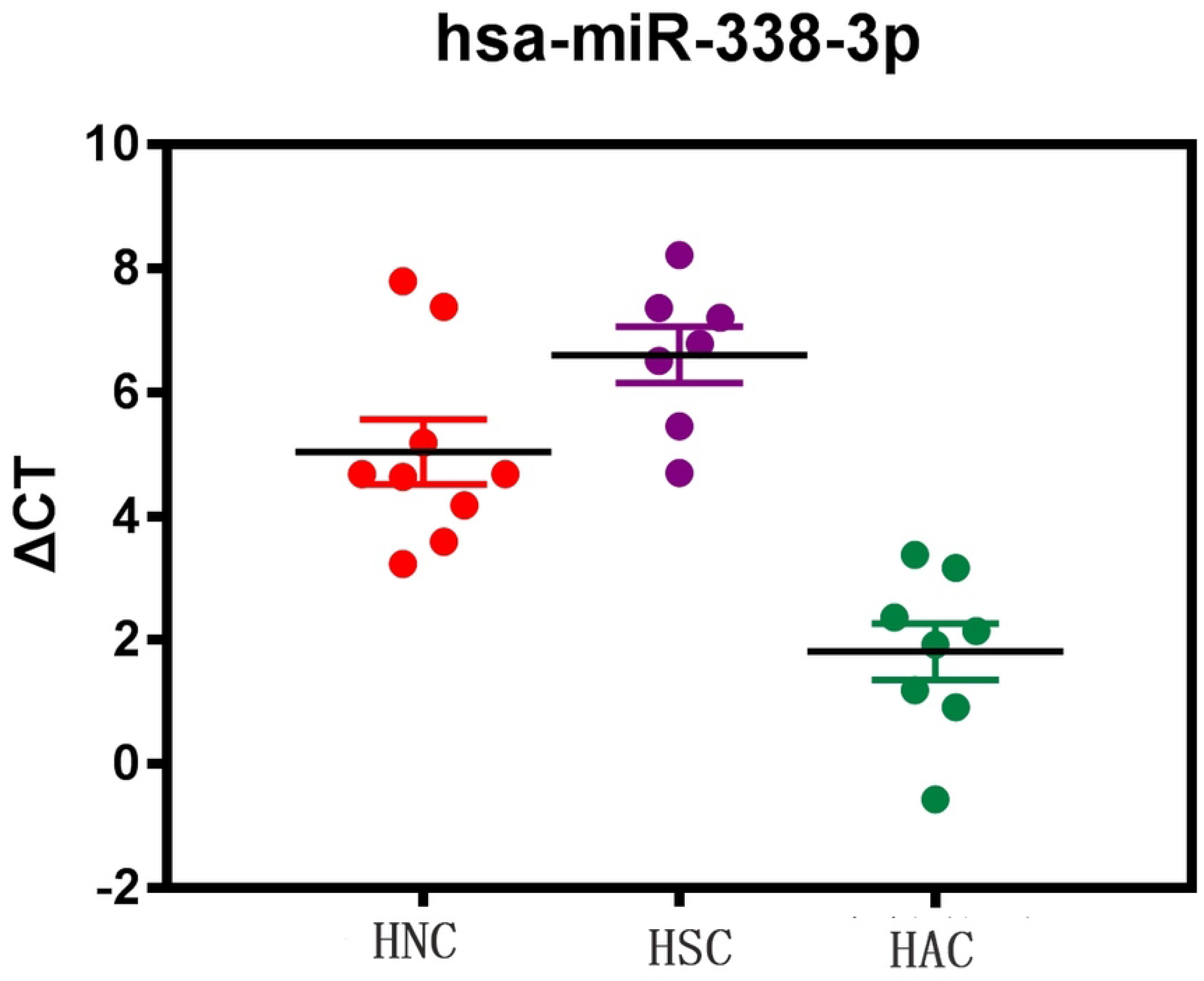
Expression of hsa-miR-338-3p among the three groups. ΔCt = Ct (gene of interest) – Ct (housekeeping gene); a higher ΔCt value indicates lower gene expression. hsa-miR-338-3p is downregulated in HSC group and upregulated in HAC group.

**Table 3.** Expression profiles of all the DEMs in the three groups

|  | HSC | HNC | HAC |
| --- | --- | --- | --- |
| hsa-miR-338-3p | ↓ | ↑ | ↑↑ |
| hsa-miR-145-5p | ↓ | = | ↑ |
| hsa-miR-183-3p | = | ↓ | ↑ |
| hsa-miR-210-3p | = | ↓ | ↑ |
| hsa-miR-490-3p | ↓ | ↑ | = |
| hsa-miR-153-3p | ↓ | ↑ | = |
| hsa-miR-489-3p | ↓ | ↑ | = |
| hsa-miR-9-5p | ↓ | ↑ | = |
| hsa-miR-212-3p | ↓ | ↑ | = |
| hsa-miR-25-5p | ↓ | ↑ | = |
| hsa-miR-127-3p | ↓ | ↑ | = |
| hsa-miR-181d-5p | ↓ | ↑ | = |
| hsa-miR-181c-3p | ↓ | ↑ | = |
| hsa-miR-9-3p | ↓ | ↑ | = |
| hsa-miR-328-3p | ↓ | ↑ | = |
| hsa-miR-608 | ↓ | ↑ | = |
| hsa-miR-205-3p | ↓ | ↑ | = |
| hsa-miR-511-5p | ↓ | ↑ | = |
| hsa-miR-129-2-3p | ↓ | ↑ | = |
| hsa-miR-338-3p | ↓ | ↑ | = |
| hsa-miR-191-3p | ↓ | ↑ | = |
| hsa-miR-431-5p | ↓ | ↑ | = |
| hsa-miR-124-5p | ↓ | ↑ | = |
| hsa-miR-142-5p | ↓ | ↑ | = |
| hsa-miR-383-5p | ↓ | ↑ | = |
| hsa-miR-187-5p | ↓ | ↑ | = |
| hsa-miR-302a-5p | ↓ | ↑ | = |
| hsa-miR-320a | = | ↑ | - |
| hsa-miR-326 | = | ↓ | ↑ |
| hsa-miR-187-3p | = | ↓ | ↑ |
| hsa-miR-770-5p | = | ↓ | ↑ |
| hsa-miR-574-3p | ↑ | = | ↓ |
| hsa-miR-382-5p | ↑ | = | ↓ |

Note: ↑ indicates upregulated expression level of the miRNA; ↑↑ indicates highly upregulated expression level of the miRNA; = indicates no significant difference was found in the expression of the miRNA compared to any group.
hsa, Homo sapiens; miR, microRNA.

### Predictive Potential of Hsa-miR-338-3p

We used receiver operating characteristic (ROC) analysis to assess the predictive potential of serum hsa-miR-338-3p levels. The area under the curve (AUC) indicated that the serum hsa-miR-338-3p expression level could potentially distinguish HSC group from the other two groups (AUC=0.908, p<0.01, Figure 4a), HAC group from the other two groups (AUC=0.992, p<0.01, Figure 4b) and HNC group from the other two groups (AUC=0.607, p<0.015, Figure 4c).

**Figure 4.**
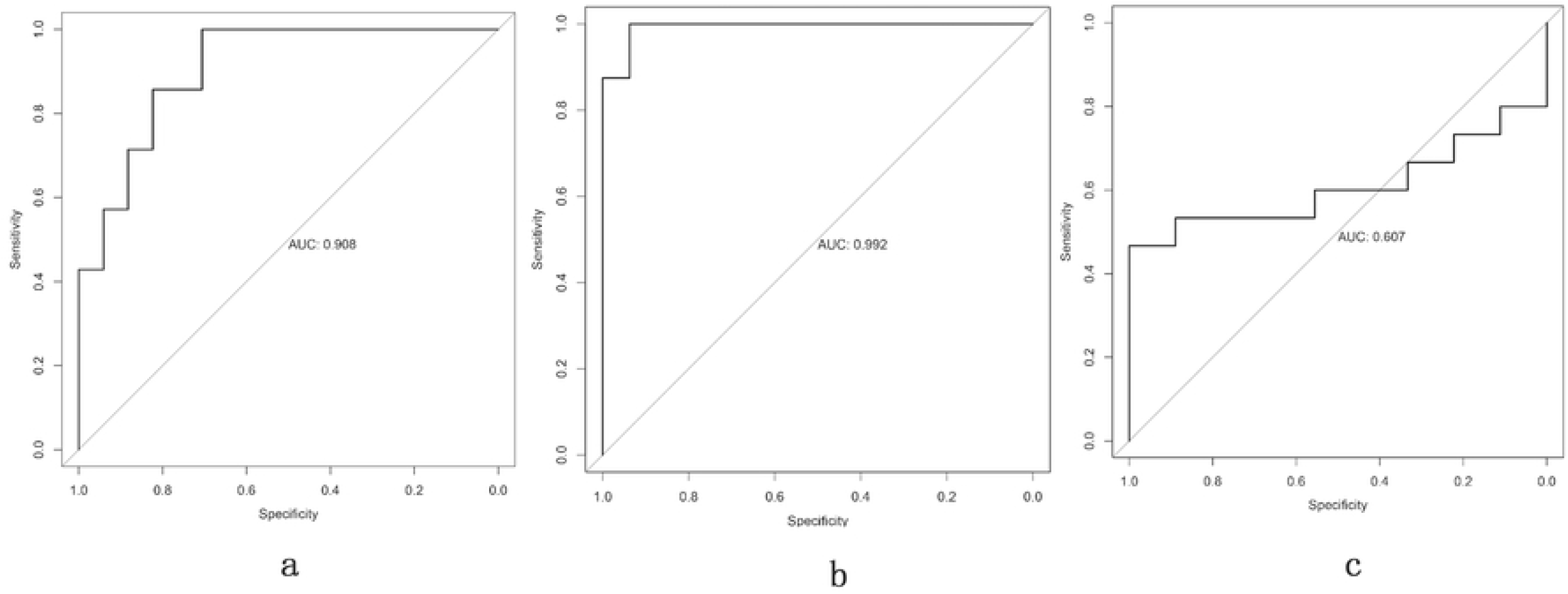
ROC analysis for serum hsa-miR-338-3p. The figure shows that serum hsa-miR-338-3p expression could potentially distinguish HSC group from the other two groups (AUC=0.908, *p*<0.01, Figure 4a), HAC group from the other two groups (AUC=0.992 *p*<0.01, Figure 4b) and HNC group from the other two groups (AUC=0.607 *p*<0.015, Figure 4c).

### Prediction of DEM Target Genes and Functional Analysis

A total of 198 targets were predicted for the common DEMs among the three groups (Table S4). Next, we annotated the biochemical signaling pathways related to the predicted miRNA targets by subjecting the predicted target genes to the KEGG pathway database (http://www.kegg.jp/). Based on the KEGG pathway enrichment analysis, we found that the predicted target genes were associated with numerous glucose/lipid metabolism-related signaling pathways and inflammation-related signaling pathways, including hsa04911: insulin secretion, hsa00564: glycerophospholipid metabolism, hsa04750: inflammatory mediator regulation of TRP channels and hsa04014: ras signaling pathway. There were 11 enriched KEGG pathways of the targets of hsa-miR-338-3p (Table 4). Among the pathways, the Ras signaling pathway is the most enriched pathway with the largest number of enriched genes (Figure S5). Accordingly, the targets of hsa-miR-338-3p, such as ETS1 and KDR, are important factors related to the regulation of vascular endothelial growth factor (VEGF) in the Ras signaling pathway. Among the miRNA targets, KCNMA1, CAMK2G, VAMP2 and CAMK2A are important factors related to glucose metabolism, including insulin secretion. Other targets, including PLD3, DGKB, AGPAT5 and PLA2G3, were enriched in glycerophospholipid metabolism (ko00564).

**Table 4.** Enriched KEGG pathways of the targets of hsa-miR-338-3p

| Pathway | Genes | *p* -value |
| --- | --- | --- |
| hsa04014: Ras signaling pathway | FGFR2, ETS1, FGF9, GRIN1, IGF1, GNG12, PLA2G3, ABL2, KDR | <0.0001 |
| hsa04720: Long-term potentiation | CAMK2G, PPP1R1A, GRIN1,<br>CAMK2A, PPP1CB | 0.002 |
| hsa05205: Proteoglycans in<br>cancer | CAMK2G, IGF1, PTCH1, MSN,<br>CAMK2A, PPP1CB, KDR | 0.0059 |
| hsa04810: Regulation of actin<br>cytoskeleton | FGFR2, FGD1, ARPC1B, FGF9, MSN,<br>GNG12, PPP1CB | 0.0074 |
| hsa04114: Oocyte meiosis | CPEB3, CAMK2G, IGF1, CAMK2A,<br>PPP1CB | 0.0131 |
| hsa05031: Amphetamine<br>addiction | CAMK2G, GRIN1, CAMK2A, PPP1CB | 0.017 |
| hsa04360: Axon guidance | SEMA6A, NRP1, PLXNB3, SRGAP3,<br>NTN1 | 0.0206 |
| hsa04911: Insulin secretion | KCNMA1, CAMK2G, VAMP2,<br>CAMK2A | 0.033 |
| hsa04713: Circadian entrainment | CAMK2G, GRIN1, GNG12, CAMK2A | 0.0436 |
| hsa00564: Glycerophospholipid<br>metabolism | PLD3, DGKB, AGPAT5, PLA2G3 | 0.0436 |
| hsa04750: Inflammatory mediator<br>regulation of TRP channels | CAMK2G, IGF1, CAMK2A, PPP1CB | 0.0471 |

Note: TRP, transient receptor potential; hsa, Homo sapiens; cAMP, cyclic adenosine monophosphate.

### Interaction Analysis

The interactions between pathways regulated by hsa-miR-338-3p, a miRNA specifically downregulated in serum from HSC group and upregulated in the serum from HAC group, were also predicted and visualized (Figure 5). In the network diagram, we observed that the pathways regulated by hsa-miR-338-3p interacted directly with each other (Figure 5). For example, the inflammatory mediator regulation of TRP channels interacts with other pathways regulated by hsa-miR-338-3p, including the Ras signaling pathway, glycerophospholipid metabolism, and insulin secretion.

**Figure 5.**
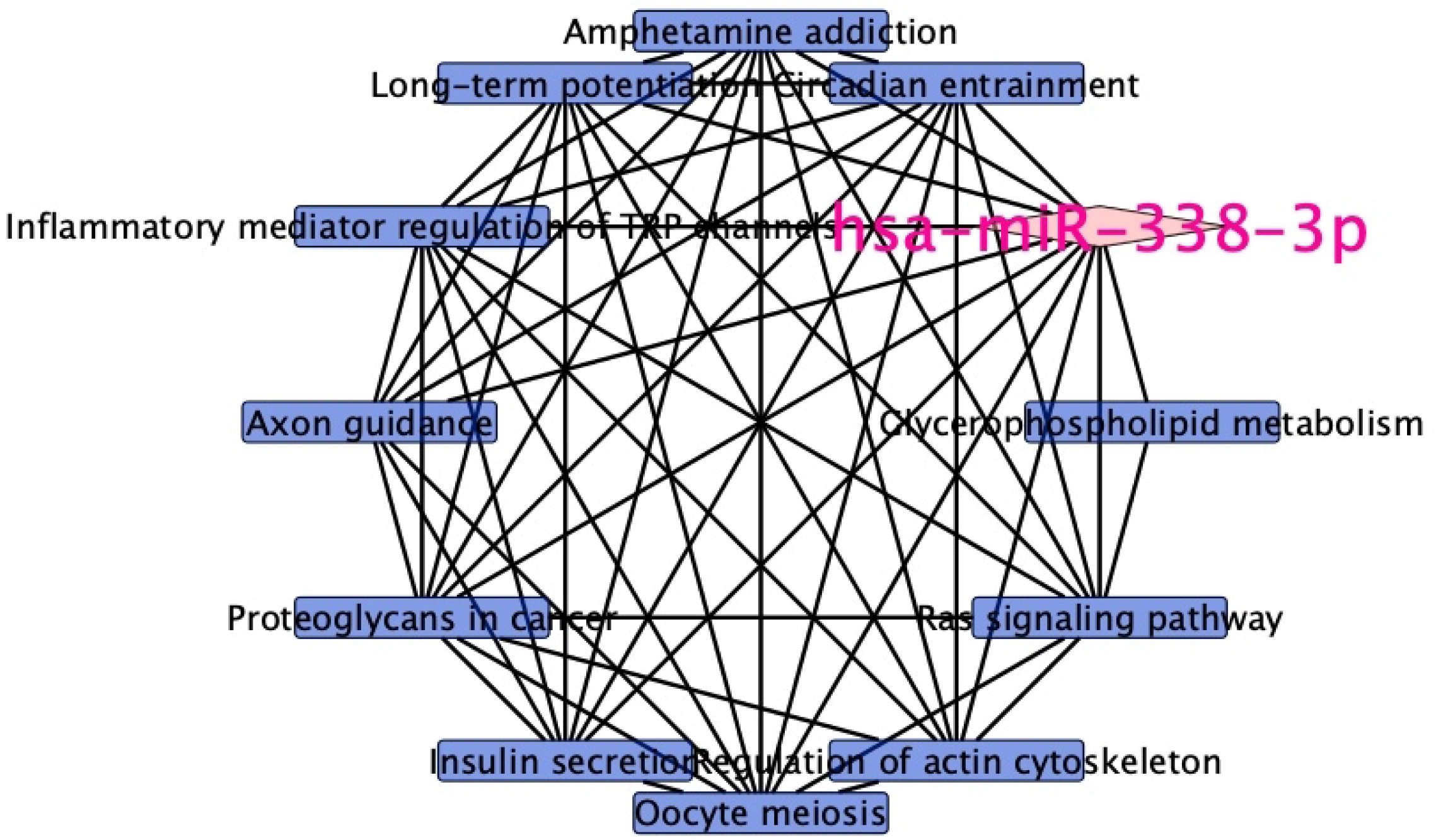
Regulatory and interaction networks of the common differentially expressed miRNA and its enriched KEGG pathways. Lines between nodes indicate interactions; nodes indicate enriched pathways.

## Discussion

Hyperlipidemia is a key factor related to atherosclerosis, cardiovascular disease, and cerebrovascular disease. TCM constitution could be an important predictor for atherosclerosis. However, the incomplete understanding of the underlying mechanisms and deficiencies in diagnosis and treatment make it difficult to prevent hyperlipidemia from progressing to atherosclerosis and vascular diseases. In the present study, using microarray analysis, we first demonstrated that global miRNA expression differences exist among hyperlipidemic patients with different TCM constitutions, and we found that hsa-miR-338-3p is a potential blood biomarker for the identification of TCM constitutions in patients with hyperlipidemia. Furthermore, strong evidence suggests that HSC is a potential early predictor of atherosclerosis complications in hyperlipidemic patients.

Our results showed that hsa-miR-338-3p was downregulated in HSC group and upregulated in HAC group compared to HNC group. Accordingly, hsa-miR-338-3p plays an important role in the negative regulation of its target genes and the involved pathways. Therefore, the target genes of hsa-miR-338-3p and the associated pathways are upregulated in HSC group and downregulated in HAC group compared to HNC group. The expression of hsa-miR-338-3p is an important factor related to atherosclerosis, cardiovascular disease, and cerebrovascular disease. Previous studies^[21, 22]^ found a strong negative correlation between hsa-miR-338-3p and high-sensitivity C-reactive protein (hs-CRP), and the low expression of hsa-miR-338-3p was a risk factor for cerebral infarction. Evidence indicates^[18, 22–24]^ that hs-CRP is an important inflammatory factor that can induce impairment and inflammation of the vascular endothelium, resulting in atherosclerosis, cardiovascular disease, and cerebrovascular disease. Therefore, hsa-miR-338-3p may be involved in the inflammatory pathway that regulates atherosclerosis and vascular disease.

To explore the molecular mechanisms underlying the atherosclerotic process in hyperlipidemic patients, we performed KEGG analysis of the target genes of hsa-miR-338-3p (Figure S5). The results demonstrated that hsa-miR-338-3p was involved in the inflammatory mediator regulation of TRP channels (TRPCs), suggesting that hsa-miR-338-3p may participate in the development of atherosclerosis. As previously reported, there are seven kinds of TRPCs. The upregulation of transient receptor potential channel 1 (TRPC1) is a general feature of smooth muscle cells in occlusive vascular disease, and TRPC1 inhibitors have potential as protective agents against human vascular failure^[25, 26]^. Another study^[27]^ found that TRPC1 maintains adherens junction plasticity by suppressing sphingosine kinase 1 expression to induce endothelial hyperpermeability and endothelial barrier destabilization, resulting in atherosclerosis formation. In addition, another study^[28]^ showed that the activation of the TRPC6 and TRPC5 channels is the key contributor to impaired endothelial healing of arterial injuries in hypercholesterolemic mice. In addition, the Ras signaling pathway appeared to be the most enriched pathway with the largest number of enriched genes. The Ras signaling pathway is well known to be involved in atherosclerosis by activating VEGF^[29, 30]^, which participates in the formation of coronary heart disease^[31]^ and cerebral infarction^[32, 33]^. In the present study, we also found that hsa-miR-338-3p may be involved in glycerophospholipid metabolism, insulin secretion and so on (Figure 5). As a previous study reported, the metabolism of glycerophospholipids plays an important role in the regulation of monocytes and macrophages^[34]^, which is a potential mechanism for atherosclerosis formation. In the network diagram (Figure 5), we can see that these pathways interact with each other, which is probably the underlying mechanism by which hsa-miR-338-3p is involved in the development of atherosclerosis. In the present study, hsa-miR-338-3p was found to be downregulated in HSC group and was inversely correlated with pathways that accelerate atherosclerosis and vascular diseases, such as the ras signaling pathway. Furthermore, we also determined that HSC group tend to develop diabetes and hyperuricemia, which are risk factors for atherosclerosis and cardiovascular and cerebrovascular diseases^[35–38]^. Thus, we hypothesized that HSC group may have a high risk for CHD and cerebrovascular disease among the hyperlipidemic population.

## Conclusion

This study uncovers that hsa-miR-338-3p is the biomarker of the TCM constitution in hyperlipidemic patients. Thus, identification of TCM constitution is no longer the privilege to TCM doctors and all clinicians will be skilled in identification of TCM, which will make TCM popularization and application. Furthermore, the present study reveals that HSC classification is a high-risk factor for the onset of atherosclerosis complications. So, intervention measures including some specific diet, exercise, herb drugs, and acupuncture will help prevent the disease by correcting patients’ constitution, which will provide better outcomes in hyperlipidemic population.

## Abbreviations

TCM: traditional Chinese medicine
DEMs: microRNAs
DEMs: differentially expressed miRNAs
HAC: hyperlipidemic patients with asthenic constitution
HSC: hyperlipidemic patients with strong constitution
HNC: 10 hyperlipidemic patients with normal constitution
ROC: receiver operating characteristic
LDL-C: low-density lipoprotein cholesterol
CHD: coronary heart disease
HLP: hyperlipidemia
TC: total cholesterol
TG: triglyceride
nonHDL-C: nonhigh-density lipoprotein cholesterol
HDL-C: high-density lipoprotein cholesterol
CT: threshold cycle
SEM: standard error of the mean
GO: Gene Ontology
KEGG: Kyoto Encyclopedia of Genes and Genomes
PCA: principal component analysis
BMI: body mass index
FT3: free triiodothyronine
FT4: free thyroxine
PCT: procalcitonin
WBC: white blood cell count
RBC: red blood cell count
PLT: platelet count
TSH: thyroid stimulating hormone
TSTO: testosterone
Hs-CRP: high-sensitivity C-reactive protein
eGFR: glomerular filtration rate
Hb: hemoglobin
UA: uric acid
GLU: glucose
hsa: Homo sapiens
miR: microRNA
ROC: receiver operating characteristic
AUC: area under the curve
VEGF: vascular endothelial growth factor
TRP: transient receptor potential
cAMP: cyclic adenosine monophosphate

## Availability of Data and Materials

The datasets generated and analyzed during the current study are available in the supplementary material.

## Compliance with ethics guidelines

All patients provided written informed consent to participate in the study. All procedures performed in studies involving human participants were in accordance with the ethical standards of the institutional and/or national research committee and with the 1964 Helsinki declaration and its later amendments or comparable ethical standards. Ethical approval was obtained from the Ethics Committee of Guangdong Provincial Hospital of Chinese Medicine (B2017-150-01).

## Authors’ contributions

Liling Zeng: conceived and designed the study and methodology, performed the investigation, experiments, data curation, and data analysis, wrote the original draft, and reviewed and edited the manuscript. Zhimin Yang: Conceptualization, funding acquisition, and reviewed and edited the manuscript. Fuping Xu: Conceptualization, data curation, formal analysis, funding acquisition, methodology, project administration, supervision, and reviewed and edited the manuscript. Qixin Zhang and Chen Sun: investigation and methodology. Jiamin Yuan: investigation and methodology. Fei Tan: investigation and methodology. Yanhua Wu: investigation and methodology.

## Acknowledgments

We would like to thank the subjects who participated in this study. The experiment was conducted at the National Engineering Center for Biochip at Shanghai and the State Key Laboratory of Dampness Syndrome of Chinese Medicine, China. The authors are grateful to Dr. Yao Qi for their technical support and assistance.

## References

[1] Ogura M, Harada-Shiba M, Masuda D, et al. Factors Associated with Carotid Atherosclerosis and Achilles Tendon Thickness in Japanese Patients with Familial Hypercholesterolemia: A Subanalysis of the Familial Hypercholesterolemia Expert Forum (FAME) Study. J Atheroscler Thromb. 2022. 29(6): 906–922.

[2] Langlois MR, Nordestgaard BG. Which Lipids Should Be Analyzed for Diagnostic Workup and Follow-up of Patients with Hyperlipidemias. Curr Cardiol Rep. 2018. 20(10): 88.

[3] Jayashree S, Arindam M, Vijay KV. Genetic epidemiology of coronary artery disease: an Asian Indian perspective. J Genet. 2015. 94(3): 539–49.

[4] Domanski MJ, Fuster V, Diaz-Mitoma F, et al. Next Steps in Primary Prevention of Coronary Heart Disease: Rationale for and Design of the ECAD Trial. J Am Coll Cardiol. 2015. 66(16): 1828–1836.

[5] Robinson JG. Is a statin as part of a polypill the answer. Curr Atheroscler Rep. 2009. 11(1): 15–22.

[6] Sandesara PB, Virani SS, Fazio S, Shapiro MD. The Forgotten Lipids: Triglycerides, Remnant Cholesterol, and Atherosclerotic Cardiovascular Disease Risk. Endocr Rev. 2019. 40(2): 537–557.

[7] Silverman MG, Ference BA, Im K, et al. Association Between Lowering LDL-C and Cardiovascular Risk Reduction Among Different Therapeutic Interventions: A Systematic Review and Meta-analysis. JAMA. 2016. 316(12): 1289–97.

[8] Simoes RG, 0000-0003-4989-4087 AO, Bernardes C, et al. Polymorphism in Simvastatin: Twinning, Disorder, and Enantiotropic Phase Transitions. Mol Pharm. 2018. 15(11): 5349–5360.

[9] Talic S, Marquina C, Zomer E, et al. Attainment of low-density lipoprotein cholesterol goals in statin treated patients: Real-world evidence from Australia. Curr Probl Cardiol. 2022. 47(7): 101068.

[10] Meurer L, Cohen SM. Drug-Induced Liver Injury from Statins. Clin Liver Dis. 2020. 24(1): 107–119.

[11] Ahmadi Y, Mahmoudi N, Yousefi B, Karimian A. The effects of statins with a high hepatoselectivity rank on the extra-hepatic tissues; New functions for statins. Pharmacol Res. 2019. 152: 104621.

[12] Huang J, Wang Y, Ying C, Liu L, Lou Z. Effects of mulberry leaf on experimental hyperlipidemia rats induced by high-fat diet. Exp Ther Med. 2018. 16(2): 547–556.

[13] Rouhi-Boroujeni H, Heidarian E, Rouhi-Boroujeni H, Khoddami M, Gharipour M, Rafieian-Kopaei M. Use of lipid-lowering medicinal herbs during pregnancy: A systematic review on safety and dosage. ARYA Atheroscler. 2017. 13(3): 135–155.

[14] Chu SM, Shih WT, Yang YH, Chen PC, Chu YH. Use of traditional Chinese medicine in patients with hyperlipidemia: A population-based study in Taiwan. J Ethnopharmacol. 2015. 168: 129–35.

[15] Lew-Ting CY, Hurwicz ML, Berkanovic E. Personal constitution and health status among Chinese elderly in Taipei and Los Angeles. Soc Sci Med. 1998. 47(6): 821–30.

[16] Huang YC, Lin CJ, Cheng SM, et al. Using Chinese Body Constitution Concepts and Measurable Variables for Assessing Risk of Coronary Artery Disease. Evid Based Complement Alternat Med. 2019. 2019: 8218013.

[17] Ma YL, 0000-0002-7683-5763 AO, Yao H, et al. Correlation between Traditional Chinese Medicine Constitution and Dyslipidemia: A Systematic Review and Meta-Analysis. Evid Based Complement Alternat Med. 2017. 2017: 1896746.

[18] Chen Y, Wu Y, Yao H, et al. miRNA Expression Profile of Saliva in Subjects of Yang Deficiency Constitution and Yin Deficiency Constitution. Cell Physiol Biochem. 2018. 49(5): 2088–2098.

[19] Joint committee for revision the Guidelines of prevention and treatment of dyslipidemia in Chinese adults. Guidelines for prevention and treatment of dyslipidemia in Chinese adults (revised in 2016). Chinese Circulation Journal. 2016. 31(10): 937–953.

[20] 中医体质分类与判定(ZYYXH/T157-2009). 世界中西医结合杂志.2009. (04): 303–304.

[21] Teng L, AUID- Oho, Meng R. Long Non-Coding RNA MALAT1 Promotes Acute Cerebral Infarction Through miRNAs-Mediated hs-CRP Regulation. J Mol Neurosci. 2019. 69(3): 494–504.

[22] Yu H, Rifai N. High-sensitivity C-reactive protein and atherosclerosis: from theory to therapy. Clin Biochem. 2000. 33(8): 601–10.

[23] Jialal I, Devaraj S. Inflammation and atherosclerosis: the value of the high-sensitivity C-reactive protein assay as a risk marker. Am J Clin Pathol. 2001. 116 Suppl: S108–15.

[24] Roh EJ, Lim JW, Ko KO, Cheon EJ. A useful predictor of early atherosclerosis in obese children: serum high-sensitivity C-reactive protein. J Korean Med Sci. 2007. 22(2): 192–7.

[25] Kumar B, Dreja K, Shah SS, et al. Upregulated TRPC1 channel in vascular injury in vivo and its role in human neointimal hyperplasia. Circ Res. 2006. 98(4): 557–63.

[26] Edwards JM, Neeb ZP, Alloosh MA, et al. Exercise training decreases store-operated Ca2+entry associated with metabolic syndrome and coronary atherosclerosis. Cardiovasc Res. 2010. 85(3): 631–40.

[27] Tauseef M, Farazuddin M, Sukriti S, et al. Transient receptor potential channel 1 maintains adherens junction plasticity by suppressing sphingosine kinase 1 expression to induce endothelial hyperpermeability. FASEB J. 2016. 30(1): 102–10.

[28] Rosenbaum MA, Chaudhuri P, Graham LM. Hypercholesterolemia inhibits re-endothelialization of arterial injuries by TRPC channel activation. J Vasc Surg. 2015. 62(4): 1040–1047.e2.

[29] Roy H, Bhardwaj S, Babu M, et al. VEGF-A, VEGF-D, VEGF receptor-1, VEGF receptor-2, NF-kappaB, and RAGE in atherosclerotic lesions of diabetic Watanabe heritable hyperlipidemic rabbits. FASEB J. 2006. 20(12): 2159–61.

[30] Guo M, Cai Y, Yao X, Li Z. Mathematical modeling of atherosclerotic plaque destabilization: Role of neovascularization and intraplaque hemorrhage. J Theor Biol. 2018. 450: 53–65.

[31] Wang Y, Huang Q, Liu J, et al. Vascular endothelial growth factor A polymorphisms are associated with increased risk of coronary heart disease: a meta-analysis. Oncotarget. 2017. 8(18): 30539–30551.

[32] Fu Y, Ni P, Ma J, et al. Polymorphisms of human vascular endothelial growth factor gene are associated with acute cerebral infarction in the Chinese population. Eur Neurol. 2011. 66(1): 47–52.

[33] Wu L, Ye Z, Pan Y, et al. Vascular endothelial growth factor aggravates cerebral ischemia and reperfusion-induced blood-brain-barrier disruption through regulating LOC102640519/HOXC13/ZO-1 signaling. Exp Cell Res. 2018. 369(2): 275–283.

[34] Nakagawa Y, Waku K. The metabolism of glycerophospholipid and its regulation in monocytes and macrophages. Prog Lipid Res. 1989. 28(3): 205–43.

[35] Perales-Torres AL CO, Castañeda Licón MT ASE, Jiménez Andrade JM. Diabetes and type of diet as determinant factor in the progression of atherosclerosis. Arch Cardiol Mex. 2016. 86(4): 326–334.

[36] Beckman JA CMA. Vascular Complications of Diabetes. Circ Res. 2016. 118(11): 1771–85.

[37] Zhao J CH, Liu N CJ, Gu Y CJ, Yang K. Role of Hyperhomocysteinemia and Hyperuricemia in Pathogenesis of Atherosclerosis. Journal of Stroke & Cerebrovascular Diseases. 2017. 26(12): 2695–2699.

[38] Lim DH LY, Park GM CSW, Kim YG LSW, et al. Serum uric acid level and subclinical coronary atherosclerosis in asymptomatic individuals: An observational cohort study. ATHEROSCLEROSIS. 2019. 288: 112–117.

